# Cell-specific nanoengineering strategy disrupts tolerogenic signaling from myeloid-derived suppressor cells to invigorate antitumor immunity in pancreatic cancer

**DOI:** 10.1101/2024.05.25.594901

**Authors:** Nilesh U. Deshpande, Anna Bianchi, Haleh Amirian, Iago De Castro Silva, Christine I. Rafie, Bapurao Surnar, Karthik Rajkumar, Ifeanyichukwu C. Ogobuiro, Manan Patel, Siddharth Mehra, Nagaraj S. Nagathihalli, Nipun B. Merchant, Shanta Dhar, Jashodeep Datta

## Abstract

Pancreatic ductal adenocarcinoma (PDAC) is characterized by intratumoral abundance of neutrophilic/polymorphonuclear myeloid-derived suppressor cells (PMN-MDSC) which inhibit T-cell function through JAK2/STAT3-regulated arginase activity. To overcome limitations of systemic inhibition of PMN-MDSCs in cancer-bearing patients—i.e., neutropenia and compensatory myelopoietic adaptations—we develop a nanoengineering strategy to target cell-specific signaling exclusively in PMN-MDSCs without provoking neutropenia. We conjugate a chemically modified small-molecule inhibitor of MDSC-surface receptor CXCR2 (AZD5069) with polyethylene glycol (PEG) and chemically graft AZD5069-PEG constructs onto amphiphilic polysaccharide derivatives to engineer CXCR2-homing nanoparticles (CXCR2-NP). Cy5.5 dye-loaded CXCR2-NP showed near-exclusive uptake in PMN-MDSCs compared with PDAC tumor-cells, cancer-associated fibroblasts, and macrophages. Encapsulation of JAK2/STAT3i Ruxolitinib (CXCR2-NP^Ruxo^) resulted in more durable attenuation in STAT3-regulated arginase activity from PMN-MDSCs and induction of cytolytic T-cell activity vs. free Ruxolitinib *in-vitro* and *in-vivo*. Cell-specific delivery of payloads via CXCR2-homing immunonanoparticles represents a novel strategy to disrupt MDSC-mediated immunosuppression and invigorate antitumor immunity in PDAC.

---

Pancreatic ductal adenocarcinoma (PDAC) is a lethal malignancy characterized by extreme therapeutic resistance and dismal survival. Beyond an undruggable genomic landscape dominated by cooperativity between *KRAS* and *TP53* mutations as well as stromal inflammation,^1, 2^ a defining hallmark of therapeutic resistance in PDAC is its tolerogenic immune microenvironment densely populated by suppressive innate immune myeloid populations— particularly neutrophilic/polymorphonuclear myeloid-derived suppressor cells (PMN-MDSC)— which infiltrate tumors frequently to drive antigen-specific and non-specific T-cell tolerance.^3^ In conjunction with a low tumor mutation burden (TMB) and non-immunogenic antigenic landscape, this PMN-MDSC mediated T-cell dysfunction renders PDAC notoriously resistant to conventional immune checkpoint (e.g., PD-1, CTLA-4) blockade as well.^4^

Previous work from our group and others have described PMN-MDSC-driven impairment of T-cell activation through a variety of mechanisms.^5, 6^ Of these, a critical pathway in PMN-MDSCs is JAK2/STAT3-regulated arginase production resulting in local depletion of arginine, which is necessary for T-cell receptor-mediated signal transduction and activation.^7, 8^ As such, previous work from our group has demonstrated that pharmacologic *neutrophil-attenuating* strategies in preclinical PDAC models not only augments adaptive immunity but also improves chemoimmunotherapy sensitivity *in vivo.*^9^ While these efforts clearly highlight the relevance of PMN-MDSC-directed therapeutics, major impediments to these approaches in human patients arise due to: (1) dose-limiting neutropenia (such as with CXCR1/2 inhibitors); (2) compensatory myelopoietic adaptations such as increase in CCR2^+^ monocyte/macrophages; and (3) off-target adverse effects.

To overcome these limitations, we engineered PMN-MDSC-directed nanoparticles to deliver payloads—such as JAK2/STAT3 inhibitors—to enhance their therapeutic index by concentrating their action in neutrophils without compromising cell viability. We then investigated if this cell-specific platform technology, which restricts targeting of STAT3-regulated arginase activity to the PMN-MDSC compartment, would improve antitumor adaptive immunity more efficiently and durably than *systemic* anti-STAT3 pharmacotherapy in preclinical PDAC models.

To identify a PMN-MDSC selective marker that would distinguish it from all other cells in the PDAC tumor microenvironment (TME), we first evaluated single-cell RNA sequencing (scRNAseq) data from human PDAC patients (n=16)^10^ and autochthonous murine models of PDAC,^11^ as well as peripheral blood mononuclear cells (PBMC) from treatment-naïve PDAC patients (n=57) treated at our center. After using differentially expressed gene signatures to attribute clusters to their putative identities in scRNAseq, *CXCR2* was near-exclusively expressed in single-cell neutrophil/PMN-MDSC transcriptomes in human PDAC tumors (**Figure 1a**); moreover, *ARG1* expression was concentrated in intratumoral PMN-MDSC sub-clusters (**Figure 1b**). Furthermore, circulating Lin^-^CD14^-^CD15^+^ PMN-MDSCs derived from PBMCs in treatment-naïve PDAC patients treated at the University of Miami (n=57) showed higher expression of *surface* CXCR2 compared with all circulating immune cell subsets (**Figure 1c**). To validate these findings in genetically engineered mouse models that phenocopy human disease, scRNAseq from *Ptf1a*^cre/+^;*LSL-Kras*^G12D/+^;*Tgfbr2*^flox/flox^ (PKT) tumors^11^ showed dominant expression of *Cxcr2* (**Figure 1d**) and *Arg1* (**Figure S1**) in neutrophilic/PMN-MDSC clusters in the TME. CXCR2 is a surface G-protein coupled receptor (GPCR) which not only transduces critical intracellular signals but is also important for lineage determination in neutrophils/PMN-MDSCs in cancer-bearing hosts^12^, making it an ideal homing target for this PMN-MDSC directed immunonanoparticle.

**Figure 1:**
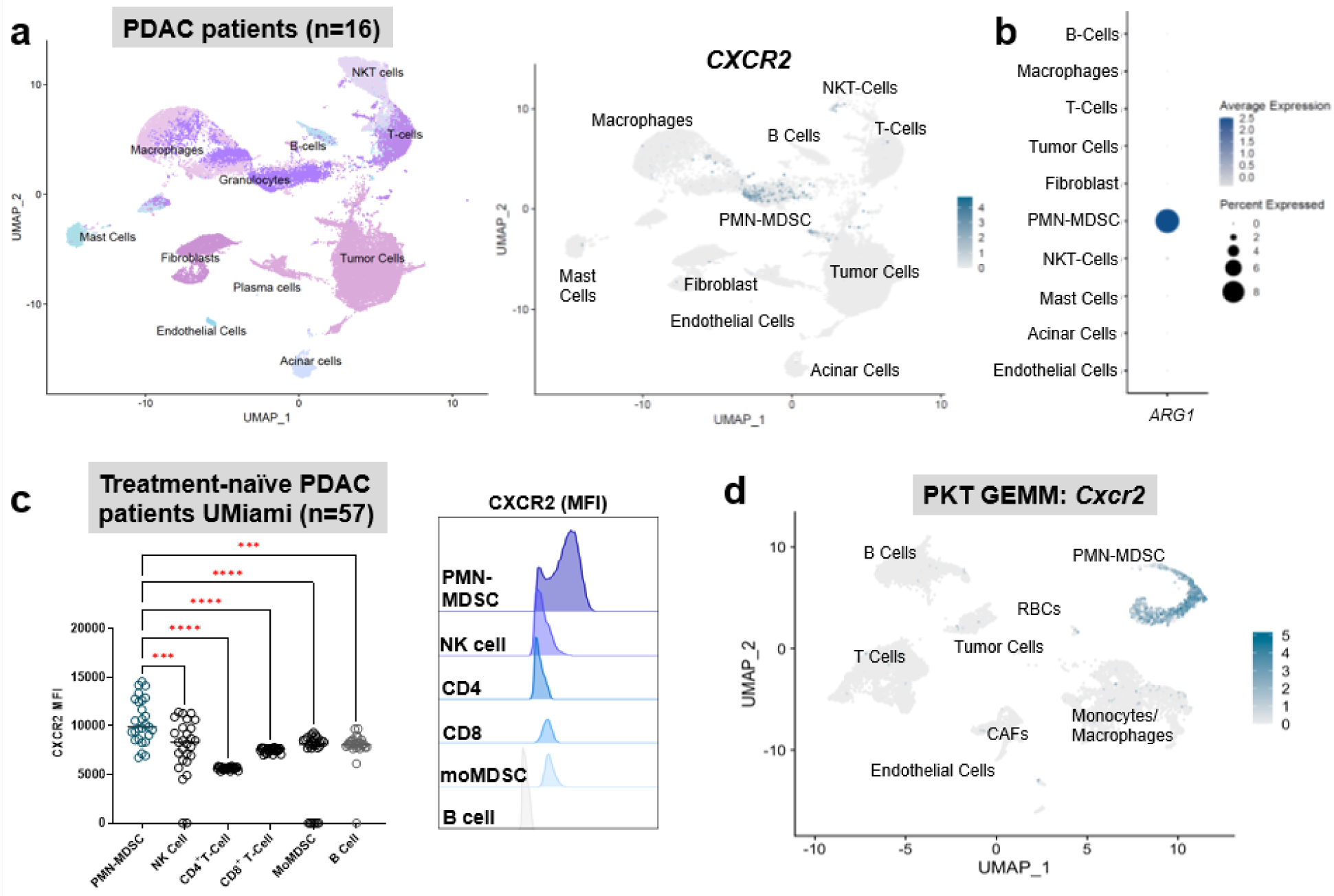
CXCR2 is an ideal PMN-MDSC-selective surface marker in human and murine PDAC. **a**, Uniform Manifold Approximation and Projection (UMAP) of single-cell RNA sequencing data (scRNAseq) from human patients with pancreatic ductal adenocarcinoma (PDAC; n=16, derived from Steele et al., *left*), with feature plot depicting *CXCR2* expression on cellular subclusters in tumor microenvironment (*right*). **b**, Bubble plot depicting arginase (*ARG1*) expression in the clusters nominated in **a**. Average expression and percent of cells expressing transcript are shown in legend. **c,** Mean fluorescence intensity (MFI) quantification of CXCR2 protein expression via flow cytometry on different circulating immune cell subsets in peripheral blood cells of treatment-naïve PDAC patients at UMiami (n= 57), depicted via bubble histogram (*left*) and pooled contour plots (*right*). **d,** UMAP of scRNAseq data from *Ptf1a*^cre/+^;*LSL-Kras*^G12D/+^;*Tgfbr2*^flox/flox^ (PKT) genetically engineered mouse model (GEMM) tumors depicting *Cxcr2* expression in cellular subclusters. ***, P<0.001; ****, P<0.0001

Our overarching goal was to engineer nanoparticles to bind specifically to CXCR2 receptors on neutrophil/PMN-MDSCs and deliver immunomodulatory payloads selectively to these cells without inducing neutropenia. As proof-of-concept, we aimed to inhibit STAT3-mediated arginase production from PMN-MDSCs, which in turns restrains arginine-mediated T-cell activation via a variety of mechanisms^13^ (**Scheme 1**).

**Scheme 1:**
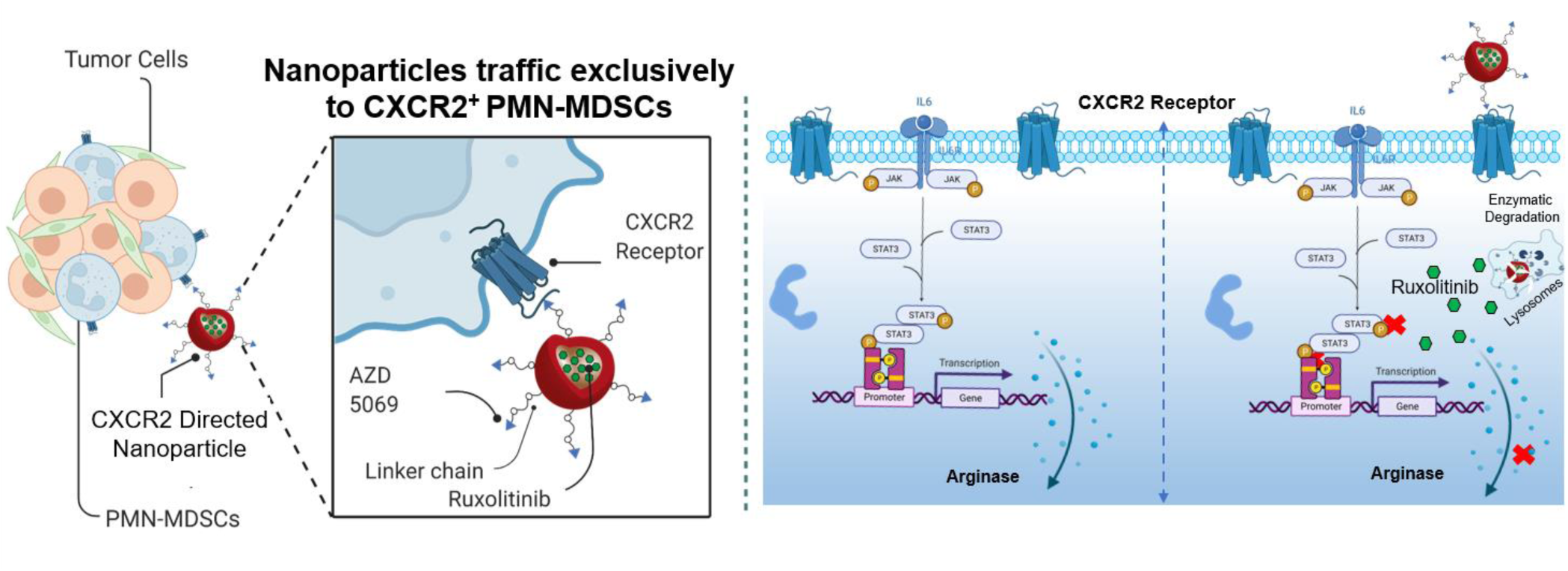
Schematic of nanoparticle design, showing how CXCR2 inhibitor AZD5069-decorated nanoparticles encapsulating hydrophobic (e.g., JAK2/STAT3 inhibitor Ruxolitinib) payloads traffic exclusively to PMN-MDSCs in the pancreatic tumor microenvironment (*left*). Upon docking at the CXCR2 receptor, this nanoparticle undergoes internalization, enzymatic degradation in lysosomes, and abrogates STAT3-mediated *Arg1* transcription and arginase activity. This reduces MDSC-instigated immunosuppression, thereby improving T-cell function.

To achieve this, we designed a biocompatible and clinical grade polysaccharide dextran-based nanocarrier by introducing appropriate amphiphilicity in the polymer backbone. Specifically, hydrophilic dextran polymer was modified with hydrophobic stearic acid and tertiary butyl succinate molecules using ester linkages that are susceptible to enzymatic degradation under lysosomal conditions (**Figure 2a**). The stearic acid and tertiary butyl succinate functional groups introduce amphiphilicity and facilitate self-assembly to generate drug-encapsulating nanoparticle. To impart CXCR2-homing capabilities to this nanoparticle backbone, we chemically modified a surface small molecule inhibitor of CXCR2 AZD5069 to conjugate this with polyethylene glycol (Boc-NH-(PEG)_36_-COOH; NMR spectroscopy characterization **Figure S2a**). The deprotection of the *Boc* group on this conjugate yielded a water-soluble derivative of AZD5069 (**Figure 2b**, **Figure S2b**), which was then chemically grafted onto the aforementioned acid-functionalized dextran derivative (**Figure 2c**) to generate CXCR2i-decorated nanoparticles (CXCR2-NP). The substitution of the H_2_N-(PEG)36-CO-AZD5069 compound on the dextran backbone was confirmed using ^1^H-NMR (**Figure S2c**).

**Figure 2:**
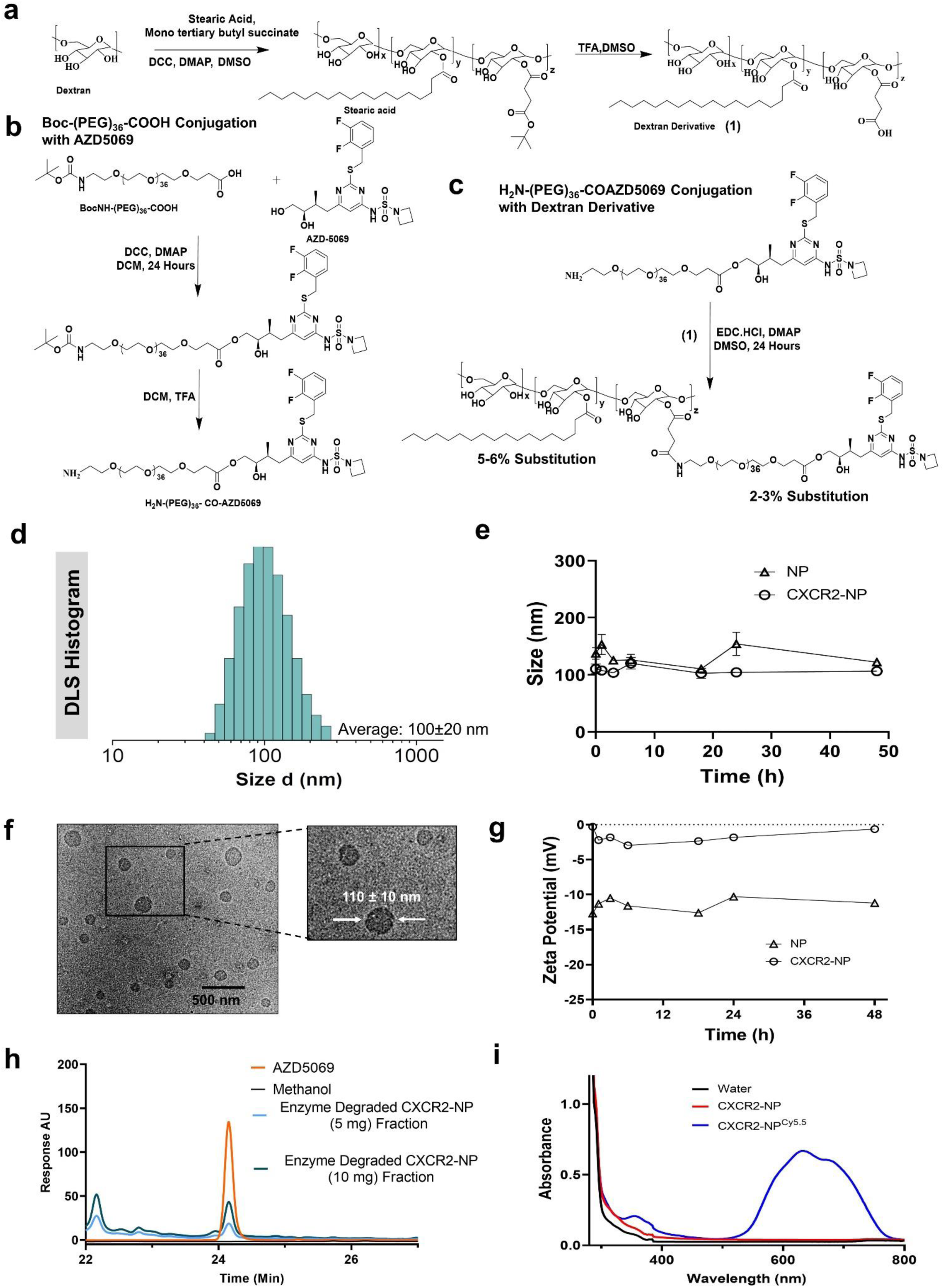
Synthetic strategy to engineer CXCR2 inhibitor AZD5069-decorated immunonanoparticles and ensuing characterization. **a,** Synthetic route showing polysaccharide dextran modification with stearic acid and tertiary butyl succinate followed by TFA deprotection to produce acid-functionalized dextran derivatives. **b,** Polyethylene glycol modification with surface CXCR2 inhibitor AZD5069 to make novel amine functionalized water soluble CXCR2i derivative. **c,** Modification of functionalized dextran derivative with newly designed CXCR2i-PEG derivative to produce CXCR2i-decorated nanoparticles (CXCR2-NP). **d,** Dynamic light scattering (DLS) histogram for CXCR2-NP showing the monomodal size distribution with average size 100±20 nm. **e,** Plot showing dynamic light scattering based stability of the blank nanoparticle and CXCR2-NP in PBS at pH 7.4 over time (hours, h). **f,** Representative transmission electron microscopy (TEM) image of CXCR2-NPs showing spherical morphology and concordant size. **g,** Plot showing zeta potential (in milliVolts, mV) of blank nanoparticles and CXCR2-NP at physiological conditions over time (hours, h). **h,** High performance liquid chromatography (HPLC) profile of free CXCR2 inhibitor AZD5069, enzyme-degraded fraction of CXCR2-NP at two concentrations (5 and 10 mg) confirming the presence of AZD5069 on the surface of CXCR2-NP. **i,** Absorbance spectrum for CXCR2-NP with or without encapsulation of Cy5.5 dye, showing expected peak at ∼683 nm in CXCR2-NP^Cy5.5^.

Self-assembly of the dextran-modified polymer in an aqueous environment resulted in monomodal particles with a Z_average_ diameter of ∼110±20 nm as determined by dynamic light scattering (DLS; **Figure 2d**). Moreover, DLS confirmed that these particles remain stable under physiological conditions up to 48 hours (**Figure 2e**). We further characterized these nano-assemblies utilizing transmission electron microscopy (TEM) to confirm morphology and hydrodynamic diameter via DLS (**Figure 2f**). Surface charge analysis by measuring zeta potential revealed a neutral surface charge on CXCR2-NPs due to conjugation of the H_2_N- (PEG)36-CO-AZD5069 moiety to dextran backbone, which exhibits surface acid functionality with ∼-15 mV charge (**Figure 2g**), indicating presence of this H_2_N-(PEG)36-CO-AZD5069 moiety on the periphery of CXCR2-NPs. High-performance liquid chromatography (HPLC) confirmed appearance of the AZD5069 peak in the organic layer extracted from enzyme-degraded CXCR2-NPs. HPLC retention time (R_t_=24.1) from CXCR2-NP aliquots exactly matched that in free AZD5069 drug aliquots, confirming retention of biophysical fidelity of CXCR2i on the surface of CXCR2-NPs (**Figure 2h**). UV-visible spectroscopy revealed encapsulation efficiency (EE) of a hydrophobic Cy5.5 dye in CXCR2-NP (CXCR2-NP^Cy5.5^) to be ∼80% (**Figure 2i**), indicating that these nanocarriers can deliver hydrophobic payloads for therapeutic purposes.

Next, we performed live-cell lysotracker imaging studies using J774 cells, which phenocopy PMN-MDSCs *in vitro* (**Figure S3a**). CXCR2-NP^Cy5.5^, structurally engineered to have an enzyme-responsive ester linkage which renders it susceptible to degradation in subcellular lysosomes (**Figure 3a**), released encapsulated Cy5.5 dye intracellularly following lysosomal degradation (**Figure 3b**) without appreciable cytotoxicity (**Figure S3b**). To examine cell-specific uptake of CXCR2-NP^Cy5.5^ in major cellular constituents present in the PDAC TME, we utilized J774 PMN-MDSCs, *K-ras^LSL.G12D/+^;p53^R172H/+^;Pdx1^Cre/+^*(KPC) tumor cells, RAW 274.1 macrophages, and KPC cancer-associated fibroblasts (CAF), confirming near-exclusive CXCR2 expression in J774 cells (**Figure 3c**). Optical imaging of these cells using confocal microscopy following incubation with CXCR2-NP^Cy5.5^ revealed incremental and selective uptake in J774 PMN-MDSC cells compared with tumor cells, CAFs, or macrophages *in vitro* at 6 hours (62% of cells/HPF), 24 hours (75%/HPF) and 48 hours (∼95%/HPF) (**Figure 3d-f**, **Figure S4a**).

**Figure 3:**
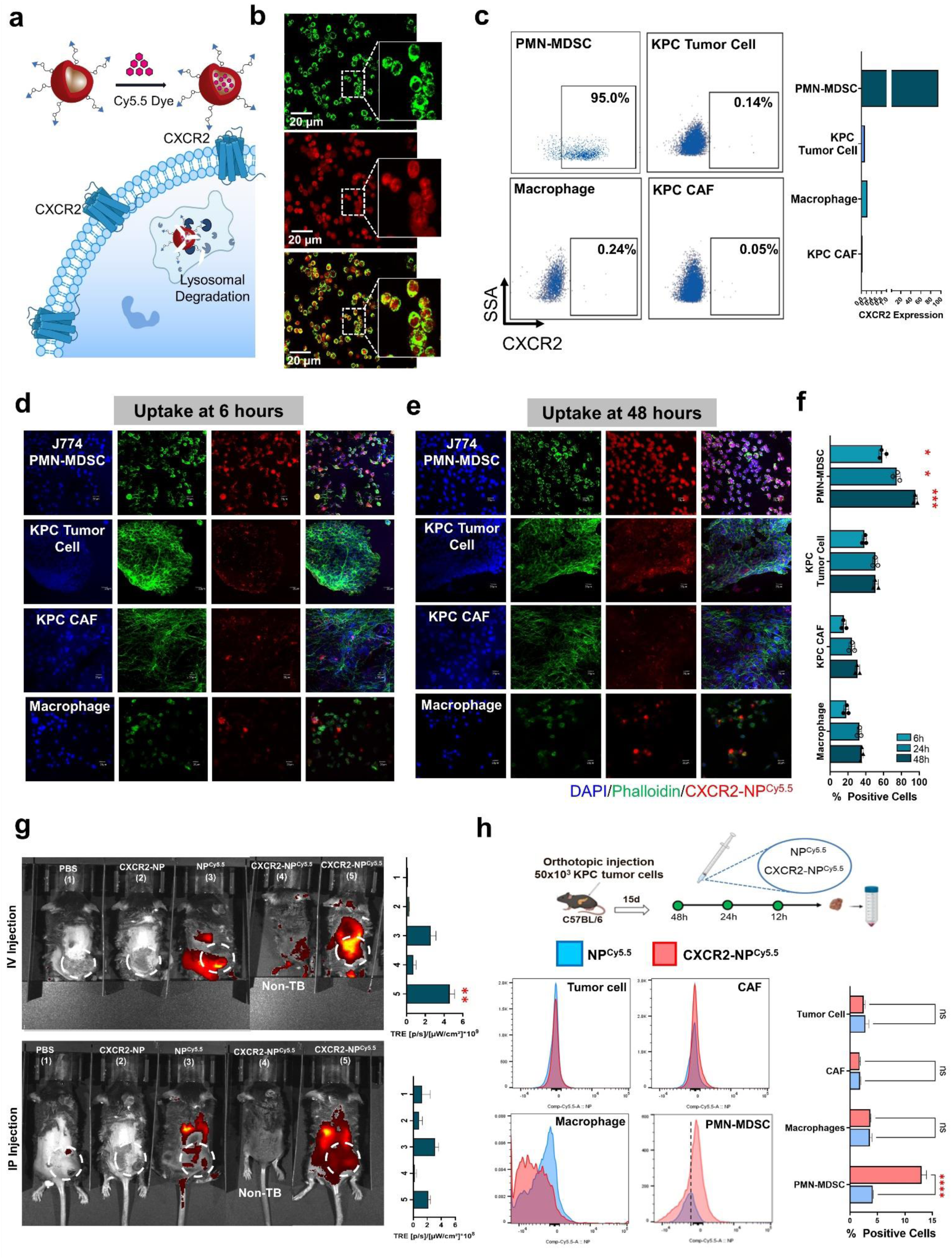
CXCR2-directed nanoparticles demonstrate preferential uptake in PDAC-associated PMN-MDSCs. **a,** Schematic showing Cy5.5 dye encapsulation in CXCR2-NP for receptor-mediated uptake of CXCR2-NP^Cy5.5^ followed by lysosomal degradation and intracellular detection. **b,** Representative confocal microscopy images in J774 cells, which phenocopy PMN-MDSCs *in vitro*, showing lysosomal co-localization (*top*) of CXCR2-NP^Cy5.5^ and release of entrapped Cy5.5 dye (*middle*, *bottom* [merged]). **c,** For subsequent *in vitro* uptake experiments, flow cytometry plots shows Cxcr2 expression on major cellular constituents in PDAC—J774 PMN-MDSC, *K-ras^LSL.G12D/+^;p53^R172H/+^;Pdx1^Cre/+^*(KPC) tumor cells, RAW 274.1 macrophages, and KPC cancer-associated fibroblasts (CAF), with aggregated relative expression of Cxcr2 across biologic replicates show in adjoining histogram. Confocal images showing cellular uptake of CXCR2-NP^Cy5.5^ in these four cell types at **d,** 6 hours post-exposure, and **e,** 48 hours post-exposure *in vitro*. **f,** Bar plot showing percent positive area for Cy5.5 positive cells at 6, 24, and 48 hour time points in these cell types, indicating progressive and near-exclusive uptake of CXCR2-NP^Cy5.5^ in PMN-MDSCs. **g,** Representative IVIS images in wildtype C57Bl/6 mice bearing KPC flank tumors (except non-tumor bearing [TB] mice in **group 4**) injected with PBS (**group 1**), non-dye loaded CXCR2-NP (**group 2**), non CXCR2i-decorated NP^Cy5.5^ (**group 3**) or CXCR2-NP^Cy5.5^ (**groups 4** [non-TB] and **5** [tumor-bearing]) by either intravenous (IV; *top*) or intraperitoneal (IP; *bottom*) administration. Adjacent bar plots show quantification of peritumoral biodistribution of trafficked Cy5.5 dye via total radiant efficiency (TRE) in IVIS (n=5/group); **h,** Cellular uptake of flow cytometry-detected Cy5.5 dye in tumor cells (EpCAM^+^), CAFs (PDPN^+^), macrophages (F4/80^+^), and PMN-MDSCs (Ly6G^+^) isolated from orthotopic KPC tumors in C57Bl/6 mice 48 hours following three IV injections of either NP^Cy5.5^ or CXCR2-NP^Cy5.5^ (shown in schematic). Representative Mean Fluorescence Intensity plots are shown, and adjacent bar plots quantify relative Cy5.5 dye detection in these cells across biologic replicates (n=5/group). *, P<0.05; **, P<0.01; ***, P<0.001; ****, P<0.0001

Next, to determine the route for *in vivo* administration that would allow optimal delivery of CXCR2-NPs to the PDAC TME, we utilized IVIS imaging in subcutaneous KPC tumor-bearing syngeneic C57/BL6 mice following either intravenous or intraperitoneal injection of vehicle control, unloaded CXCR2-NP, non-AZD5069 decorated control-NP^Cy5.5^, or CXCR2-NP^Cy5.5^ constructs. Although control-NP^Cy5.5^ and CXCR2-NP^Cy5.5^ trafficked to tumor sites regardless of intravenous or intraperitoneal administration, *intravenous* injection of CXCR2-NP^Cy5.5^ demonstrated the strongest peritumoral biodistribution (P<0.01; **Figure 3g**). In orthotopic KPC tumor cell-CAF co-injection PDAC models, CXCR2-NP^Cy5.5^ but not control-NP^Cy5.5^ constructs preferentially concentrated in intratumoral Ly6G^+^F4/80^-^ PMN-MDSCs compared with F4/80^+^ macrophages, EpCAM^+^ tumor cells, PDPN^+^ CAFs (**Figure 3h**), F4/80^-^Ly6C^+^ monocytic MDSCs, CD3^+^ T-cells, and Cd11c^+^ dendritic cells (**Figure S4b**) by flow cytometry. Therefore, we utilized intravenous delivery for all subsequent *in vivo* experiments.

Having shown that CXCR2-NPs traffic reliably to PMN-MDSCs *in vitro* and *in vivo*, we next focused on packaging hydrophobic JAK2/STAT3i Ruxolitinib using the principles of aqueous self-assembly (**Figure 4a**). In pathologically activated neutrophils, it is well known that JAK2/STAT3 signaling mediates arginase production, which causes local depletion of arginine which in turn impairs cytolytic activity of T-cells via a variety of mechanisms.^13^ Upon encapsulation of Ruxolitinib in CXCR2-NPs (CXCR2-NP^Ruxo^), we confirmed that the biophysical properties of the previously characterized nanoparticles, including spherical conformation and diameter (**Figure 4b**), remain unaltered. The calculated EE via absorbance spectroscopy for CXCR2-NP^Ruxo^ was ∼80%, near-identical to EE of CXCR2-NP^Cy5.5^ (**Figure 4c**). Next, degradation dynamics of CXCR2-NP^Ruxo^ incubated with porcine liver esterase enzyme at 37^0^C in mini-dialysis units demonstrated that 90% of encapsulated Ruxolitinib is released within 48 hours (**Figure 4d**) with only 8-10% of encapsulated drug leaching out in the absence of esterase (**Figure 4e**). Together, these assays suggested reliable time-controlled release of STAT3i upon intracellular degradation of CXCR2-NPs in target cells.

**Figure 4:**
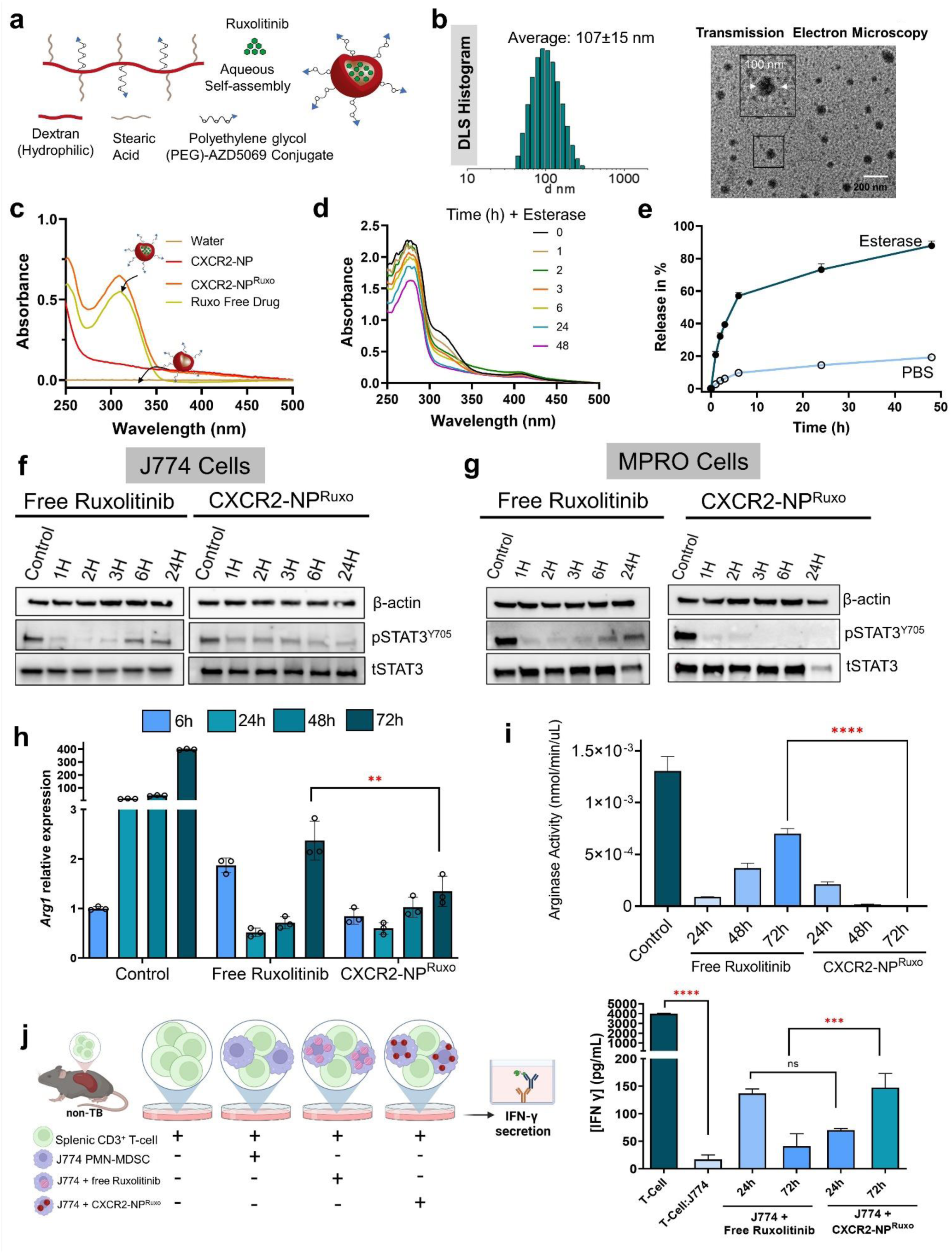
Packaging JAK2/STAT3 inhibitor in CXCR2-NP mitigates PMN-MDSC arginase activity and rescues T-cell suppression *in vitro*. **a,** Schematic showing encapsulation of JAK2/STAT3 inhibitor Roxulitinib in and self-assembly of CXCR2 inhibitor AZD5069-decorated PEG-dextran nanoparticle (CXCR2-NP^Ruxo^). **b**, Dynamic light scattering (DLS) histogram exhibiting monomodal distribution with concordant size, and transmission electron microscopy images confirming spherical morphology of nanoobjects. **c,** Absorbance spectrum of free Ruxolitinib drug, non-drug encapsulated CXCR2-NP and CXCR2-NP^Ruxo^ nanoparticles. **d,** Absorbance spectrum confirming esterase enzyme-assisted degradation of CXCR2-NP^Ruxo^ and ensuing release of payload. **e,** Plot showing percent drug release over time (in hours, h) following exposure to esterase enzyme vs. PBS, confirming slow sustained release of 90% of encapsulated Ruxolitinib over 48 hours. Western blot showing levels of pSTAT3^Y705^, total (t)STAT3, and β-actin over 24 hours following single treatment with free Ruxolitinib drug or CXCR2-NP^Ruxo^ nanoparticles in **f,** J774 cells or **g,** MPRO cells, which both phenocopy PMN-MDSCs *in vitro*. Bar plot showing relative **h,** gene expression of *Arg1* via RT-PCR and **i,** arginase functional activity via luminescence assays in J774 cells following single treatment with isotype control, free Roxulitinib, and CXCR2-NP^Ruxo^ nanoparticles up to 72 hours. **j,** Schematic summarizing experimental design, and adjacent bar plots quantifiying IFN-γ secretion (in pg/mL) from splenic CD3^+^ T-cells (derived from non-tumor bearing mice) co-cultured with either J774 cells alone, or J774 cells pre-conditioned with free Ruxolitinib drug or CXCR2-NP^Ruxo^ nanoparticles at 24 and 72 hours. Conditioned media from T-cell and J774 cell co-cultures were collected 24 hours post-incubation and subjected to IFN-γ ELISA. **, P<0.01; ***, P<0.001; ****, P<0.0001

To examine this further at a functional level, we treated two cell lines that phenocopy PMN-MDSCs *in vitro*—J774 and MPRO^14, 15^—with either free Ruxolitinib or CXCR2-NP^Ruxo^ and tested the efficacy and durability of STAT3 inhibition in neutrophils. Both free Ruxolitinib and CXCR2-NP^Ruxo^ were effective at mitigating pSTAT3^Y705^ expression via western blotting at early (1-3 hour) timepoints in J774 and MPRO cells (**Figure 4f&g**). However, while pSTAT3 expression was reactivated between 6-24 hours in both J774 and MPRO cells following free Ruxolitinib treatment, pSTAT3 expression remained durably inhibited when Ruxolitinib was delivered in CXCR2-NP^Ruxo^ in both J774 (**Figure 4f**) and MPRO (**Figure 4g**) cells.

We next sought to determine if sustained pSTAT3 inhibition in PMN-MDSCs via CXCR2-NP^Ruxo^ treatment would result in durable repression of arginase enzymatic activity and disinhibition of T-cell suppression in MDSC:T-cell co-cultures. Treatment of J774 cells with CXCR2-NP^Ruxo^ resulted in durable inhibition of both *Arg1* expression via qPCR (P=0.002; **Figure 4g**) and abolition of enzymatic arginase activity via luminescence assays (P<0.001; **Figure 4h**) compared with that observed in J774 cells treated with free Ruxolitinib. Notably, when we co-cultured J774 cells treated with either CXCR2-NP^Ruxo^ or free Ruxolitinib with CD3/CD28-stimulated T-cells from tumor-naïve C57/Bl6 mice, we observed significantly improved IFN-γ release at 72 hours when CXCR2-NP^Ruxo^-treated PMN-MDSCs were used in T-cell co-cultures compared with free Ruxolitinib-treated PMN-MDSCs (P<0.001; **Figure 4i**).

Finally, we treated orthotopic KPC tumor-bearing C57/Bl6 mice (n=10/group) with either vehicle, intraperitoneal free Ruxolitinib, non-drug loaded CXCR2-NP, or CXCR2-NP^Ruxo^ (**Figure 5a**). Importantly, we did not observe any difference in viability of splenic/circulating neutrophils across four treatment groups during (**Figure 5b**) or at endpoint analysis (**Figure S5a**), validating a cornerstone principle of our nanoengineering strategy. Tumor weights were modestly but significantly reduced in CXCR2-NP^Ruxo^-treated mice compared with vehicle, Ruxolitinib, or CXCR2-NP treatment (P<0.05 for all; **Figure 5c**). Next, to ascertain functional endpoints across treatment cohorts, we assessed pSTAT3 expression and arginase activity in *intratumoral* Ly6G^+^ PMN-MDSCs retrieved from mice in each treatment group. Expression of pSTAT3^Y705^ via western blotting (**Figure 5d**) and arginase functional activity via luminescence assays (**Figure 5e**) were significantly reduced in intratumoral PMN-MDSCs isolated from CXCR2-NP^Ruxo^-treated mice compared with vehicle, Ruxolitinib, or CXCR2-NP-treated mice. Although the overall intratumoral infiltration of CD4^+^ (**Figure S5b**) and CD8^+^ T-cells (**Figure 5f**) was not significantly different between cohorts, tumors in CXCR2-NP^Ruxo^-treated mice demonstrated significantly increased proportions of CD44^+^CD62L^-^effector, CD44^+^CD62L^+^ central memory, and degranulating CD107^+^ CD8^+^ T-cells compared with vehicle, Ruxolitinib, or CXCR2-NP treatment via flow cytometry (**Figure 5f**). Importantly, intratumoral populations of IFN-γ^+^ CD8^+^ T-cells was significantly increased in CXCR2-NP^Ruxo^-treated vs. free Ruxolitinib-treated mice (**Figure 5g**).

**Figure 5:**
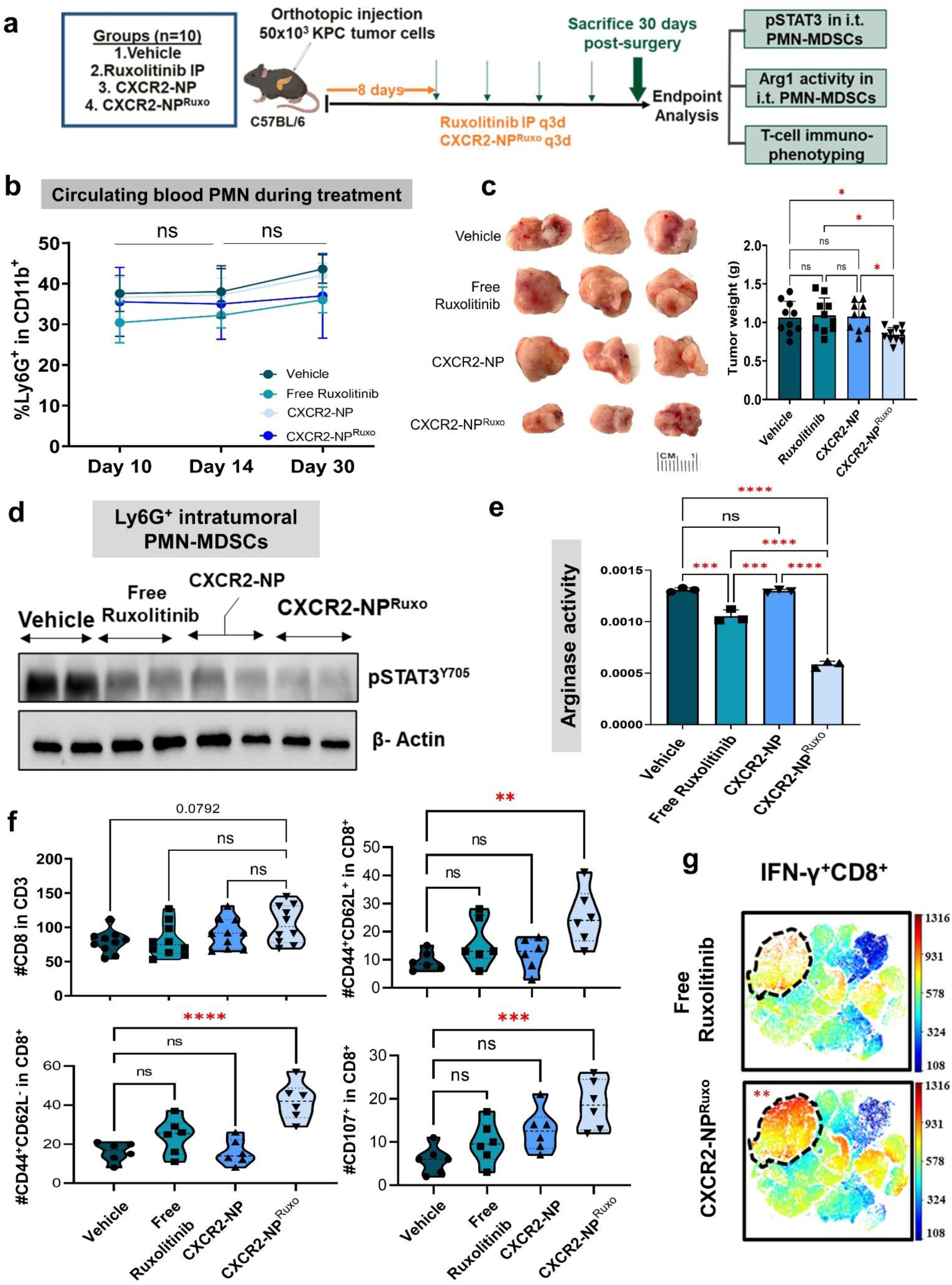
Deploying JAK2/STAT3 inhibitor in CXCR2-nanoparticles mitigates MDSC-specific arginase activity and invigorates T-cell activity in preclinical models of pancreatic cancer. **a,** Schematic of *in vivo* experiments in which mice were orthotopically injected with *K-ras^LSL.G12D/+^;p53^R172H/+^;Pdx1^Cre/+^*(KPC) tumor cells and randomized into groups (n=10 mice/group) 8 days post-surgery for treatment with vehicle control, intraperitoneal (IP) Roxulitinib, non-drug encapsulated CXCR2-NP, and CXCR2-NP^Ruxo^. Mice were sacrificed at day 30 post-surgery for endpoint analysis, as indicated. **b,** Frequencies of circulating Ly6G^+^ neutrophil/polymorphonuclear (PMN) cells (in CD11b^+^ gate) quantified in blood of mice via flow cytometric analysis at indicated time points (days, d) during treatment. **c,** Representative tumors from mice in each group (*left*) with adjacent bar plots quantifying tumor weights (in gram, g) from across treatment groups (n=10/group; *right*). **d,** pSTAT3^Y705^ protein levels by western blotting in intratumoral Ly6G^+^ PMN-MDSCs from mice in each treatment group (n=2/group shown). **e,** Arginase activity via luminescence assays in isolated intratumoral PMN-MDSCs from tumors in treated mice. **f,** Violin plots quantifying aggregated data from intratumoral T-cell immunophenotyping experiments via flow cytometry in mice across treatment groups (n=6/group). Data are shown for CD3^+^CD8^+^, CD8^+^CD44^+^CD62L^-^ (effector memory), CD8^+^CD44^+^CD62L^+^ (central memory), and CD8^+^CD107^+^ (degranulating) T-cell populations. **g,** Additonal samples from Ruxolitinib and CXCR2-NPRuxo-treated mice were fixed for intracellular staining, with viSNE plot visualizing comparison of intratumoral IFN-γ^+^ CD8^+^ T-cells from mice in these 2 groups (n=6/group). *, P<0.05; **, P<0.01; ***, P<0.001; ****, P<0.0001

The myelo-enriched immune microenvironment, and suppressive signaling from its cellular constituents PMN-MDSCs and M2-like macrophages, are key drivers of therapeutic resistance and dismal survival in PDAC.^16^ Preclinical data from our and others’ groups have demonstrated the value of disrupting PMN-MDSC-mediated signaling to overcome therapeutic resistance, yet clinical translation of neutrophil-directed *cytotoxic* strategies (e.g., CXCR1/2 inhibition) has proven challenging due to dose-limiting neutropenia and compensatory myelopoietic adaptations.^17^ We propose a PMN-MDSC-directed immunonanoengineering strategy in which CXCR2-*homing* nanoparticles deliver pharmacologic payloads with precision to PMN-MDSCs without provoking neutropenia. Utilizing proof-of-concept JAK2/STAT3 inhibition via Ruxolitinib to mitigate arginase-mediated T-cell suppression, we show not only more durable attenuation in tolerogenic STAT3-regulated arginase signaling in PMN-MDSCs but also invigoration of T-cell activity *in vitro* and *in vivo*.

Most nanotherapeutics targeting solid tumors rely heavily on *passive* accumulation in both tumor and non-tumor tissue via leaky vasculature, which renders many of these approaches non-specific. Moreover, nanoparticles directed at receptors on *cancer cells* have been disappointing due to biodistributions similar to their non-targeted counterparts^18^. In contrast, our strategy leverages tumor-associated myeloid cells—which *actively* traffic to tumor beds down tumor-derived chemokine gradients (e.g., Cxcl1-CXCR2^19^)—as endogenous cargo vectors for a diverse array of immunomodulatory payloads. This allows concentration of nanoparticle-ferried payloads at sites of active tumor-induced inflammation. Furthermore, given our growing understanding that key signaling pathways have competing and paradoxical effects in the TME depending on their *cell type of origin* (e.g., TNF signaling, NLRP3 inflammasome activation, MAP kinase activation^5, 11, 20^), an obvious advantage of our platform would be to tailor compartment-specific pharmacologic or genetic engineering approaches to thwart tolerogenic signaling while preserving immunostimulatory signaling in a cell-specific manner.

Our novel nanoengineering strategy has several other advantages, namely (a) delivery of substantially higher doses of encapsulated payload than can be delivered or tolerated systemically; (b) packaging of hydrophobic drugs or molecules that are difficult to administer intravenously; (c) sustained release of entrapped payload for more durable target inhibition in PMN-MDSCs; (d) avoiding paradoxical impact on immune-*potentiating* target signaling in disparate cell types; and (e) strong potential for clinical translation to PDAC patients given selection of FDA-approved polysaccharide dextran as the backbone for our immunonanoparticle design. Indeed, we have initiated a translatable scale-up process using previously optimized manufacturing procedures^21^ to approach regulatory path for this technology, with the ultimate goal of human translation via phase 1 trials in PDAC patients.

Given the paucity of actionable myeloid-directed immunotherapeutics in patients with solid tumors, our novel nanoengineering strategy holds promise to not only overcome historical drawbacks but also offer a platform to target MDSC-intrinsic pathways mediating T-cell and stromal dysregulation in PDAC. Moreover, future efforts must focus on rational combinations with tumor cell-autonomous signaling (e.g., *KRAS* inhibition) and T-cell directed strategies (e.g., PD-1/CTLA-4 inhibitors, CAR-T, T-cell engagers, etc.) to further augment the therapeutic index and improve outcomes in this devastating malignancy.

## Supporting information

Supporting Information (Methods, Suppl Figures/Tables, References)

## SUPPORTING INFORMATION

Additional experimental details, materials, and methods, Supplementary figures, including ^1^H-NMR spectra for all compounds (PDF)

## ACKNOWLEDGMENTS

Senior author JD receives research funding from Cantargia AB, Inc. (Sweden), unrelated to this work. The remainder of authors declare no competing financial interest. This work was supported by U.S. Department of Defense Idea Development Award grant # HT9425310699, Sylvester Transdisciplinary Science Pilot grant # TDSP-2023-02, and Pancreatic Cancer Action Network Career Development Award grant # 22-20-DATT (J.D.). Research reported in this publication was supported by the National Cancer Institute of the National Institutes of Health under Award Number P30CA240139. The content is solely the responsibility of the authors and does not necessarily represent the official views of the National Institutes of Health.

