## Supporting Information (Methods, Suppl Figures/Tables, References) for "Cell-specific nanoengineering strategy disrupts tolerogenic signaling from myeloid-derived suppressor cells to invigorate antitumor immunity in pancreatic cancer"

**for**

[1] Department of Surgery, University of Miami Miller School of Medicine, Miami FL
[2] Department of Biochemistry and Molecular Biology, University of Miami Miller School of Medicine, Miami FL
[3] Sylvester Pancreatic Cancer Research Institute, Sylvester Comprehensive Cancer Center, Miami FL
[4] Engineering Cancer Cures™ Program

**CONTENTS**


**1. Materials and Methods**

1.1. Synthesis of dextran-stearic acid-succinic mono tert. butyl ester derivative
1.2. Synthesis of dextran-stearic acid-succinic acid derivative.
1.3. Synthesis of Boc-N-amido-$(PEG)_{36}$-CO-AZD5069 conjugate.
1.4. Synthesis of $H_2N$-$(PEG)_{36}$-CO-AZD5069 conjugate.
1.5. Synthesis of dextran-stearic acid-succinic acid- $(PEG)_{36}$-AZD5069 derivative.
1.6. Biophysical characterization of CXCR2-homing nanoparticle.
1.7. Single-cell RNA sequencing (scRNAseq) analysis.
1.8. Cluster identification and gene annotation in scRNAseq dataset.
1.9. Cell Lines.
1.10. Arginase Activity Luminescence Assay.
1.11. Enzyme-linked immunosorbent assay (ELISA).
1.12. Western Blotting.
1.13. Quantitative Polymerase Chain Reaction (qPCR).
1.14. Orthotopic and subcutaneous mouse models.
1.15. Tissue Processing.
1.16. Flow Cytometry.
1.17. IVIS imaging experiments.
1.18. T-cell suppression assays.

**2. Supplementary Tables**

2.1. Table S1
2.2. Table S2

**3. Supplementary Figures**

3.1. Figure S1
3.2. Figure S2
3.3. Figure S3
3.4. Figure S4
3.5. Figure S5

**4. References**

### 1. MATERIALS AND METHODS

***1.1. Synthesis of dextran-stearic acid-succinic mono tert. butyl ester derivative (compound 1).*** Dextran ($M_w$= 6,000, 1.0 g, 6.2 mmol of anhydroglucose unit), stearic acid (0.2 g, 0.80 mmol) and tert. butyl succinate (0.2 g, 1.15 mmol) were dissolved in anhydrous DMSO:DMF:DCM (6mL+ 4mL+ 2mL) solvent mixture. To the above content DMAP (0.2 g, 1.63 mmol), EDC.HCl (0.24 g, 1.26 mmol) and NHS (0.14 g, 1.21 mmol) were added and the reaction mixture was stirred for 24 hour at room temperature. The reaction mixture was dialyzed against water using 3.5KDa MWCO tubing to remove reaction byproducts, frozen and lyophilized. The obtained solid / thick viscous liquid was precipitated by adding into methanol and acetone solvent mixture and solid product was separated by centrifugation. This solid product was washed several times with methanol: acetone solvent mixture and obtained solid was dissolved in water, frozen and lyophilized to get white powder as a pure product (**Figure 2**, **Figure S2a**). Yield = 1.2 g (36 %). $^1$H-NMR (400 MHz, $d_6$-DMSO) δ ppm: 4.47, 4.82, 4.88 ppm (s, hydroxyl of dextran) 4.63 ppm (s, dextran anomeric proton), 3.14-3.69 ppm (dextran glucosidic protons), 2.6 ppm (m,4H) methylene protons of SA, 1.48 ppm,1.39 ppm (s, 9H) tert. butyl, 1.25-0.85 ppm (aliphatic protons).

***1.2. Synthesis of dextran-stearic acid-succinic acid derivative.*** Compound **(1)** (0.7 g, 1.27 mmol) was dissolved in 2.0 mL of DMSO and 200μL of trifluoro acetic acid (TFA) was added to it. The resulting reaction mixture was stirred for one hour at room temperature. The reaction mixture was then dialyzed against water using 3.5KDa MWCO tubing to remove reaction byproducts, frozen and lyophilized. The obtained solid / thick viscous liquid was precipitated by adding into methanol and acetone solvent mixture and the solid product was separated by centrifugation. This solid product was then washed several times with methanol: acetone solvent mixture and finally a solid fraction was obtained and dissolved in water, frozen and lyophilized to obtain a pure product (**Figure 2**, **Figure S2b**). Yield = 0.45 g (78 %). $^1$H-NMR (500 MHz, $d_6$-DMSO) δ ppm: 4.47, 4.82, 4.88 ppm (s, hydroxyl of dextran) 4.63 ppm (s, dextran anomeric proton), 3.14-3.69 ppm (dextran glucosidic protons), 2.6 ppm (m,4H) methylene protons of SA, 1.48 ppm, 1.25-0.85 ppm (aliphatic protons).

***1.3. Synthesis of Boc-N-amido-(PEG)$_{36}$-CO-AZD5069 conjugate.*** Boc-N-amido-(PEG)36-acid, (0.1 g, 0.056 mmol) and AZD5069, 27 mg (0.056 mmol) were dissolved in 2.0 mL of dry dichloromethane solvent and stirred for 10 minutes at room temperature. To the above reaction mixture, DCC (18 mg, 0.087 mmol) and DMAP (10 mg, 0.081 mmol) were added and the resulting solution was stirred for 24 hours at room temperature. The DCU formed was separated by filtration, DCM in the reaction mixture was evaporated in rota evaporator and the obtained viscous liquid was precipitated by adding it into cold diethyl ether. The solid product obtained was washed several times with cold diethyl ether and then poured into water to remove the unreacted AZD5069 traces. Finally, the water-soluble product was frozen and lyophilized to obtain a white solid powder as Boc-N-amido-(PEG)36-CO-AZD 5069 conjugate (**Figure 2**, **Figure S2c**). Yield: 100 mg (86%) $^1$H-NMR (500 MHz, $d_6$-DMSO) δ ppm: 7.1-7.5 (4H, AZD5069), 6.1(1H, AZD 5069), 4.44 (2H, PEG), 3.3-3.8 (165H, PEG and AZD5069), 3.0-3.1 (8H, PEG + AZD 5069), 1.35 (9H, tert butyl group Boc-N-amido-(PEG)$_{36}$ ), 1.18-1.22 (5H, from AZD5069).

***1.4. Synthesis of $H_2N$-(PEG)$_{36}$-CO-AZD5069 conjugate.*** Boc-N-amido-(PEG)36-CO-AZD5069 conjugate (80 mg, 0.038 mmol) was dissolved in 2.0 mL of dry dichloromethane solvent and stirred for 10 minutes at room temperature. Then, TFA (200 μL) was added dropwise and the resulting solution was stirred for 1 hour at room temperature, DCM in the reaction mixture was evaporated using rota evaporator and the obtained viscous liquid was precipitated by adding it into cold diethyl ether. The solid product obtained was washed several times with cold diethyl ether, separated by centrifugation, dissolved in water lyophilized to obtain viscous liquid as $H_2N$-(PEG)36-CO-AZD 5069 conjugate (**Figure 2**, **Figure S2d**). Yield: 70 mg (92%) $^1$H-NMR (500 MHz, $d_6$-DMSO) δ ppm: 7.1-7.5 (4H, AZD5069), 6.1(1H, AZD 5069), 4.44 (2H, PEG), 3.3-3.8 (165H, PEG and AZD5069), 3.0-3.1 (8H, PEG + AZD 5069), 1.18-1.22 (5H, from AZD5069).

***1.5. Synthesis of dextran-stearic acid-succinic acid- (PEG)$_{36}$-AZD5069 derivative.*** Dextran-stearic acid-succinic acid derivative (50 mg, 0.19 mmol) was dissolved in 1.0 mL of dry DMSO. To the above reaction mixture, EDC.HCl (54 mg, 0.282 mmol), DMAP (23 mg, 0.18 mmol), NHS (32mg, 0.278 mmol) were added and stirred for 30 minutes. In another vial, $H_2N$-(PEG)$_{36}$- CO-AZD5069 (20 mg, 0.010 mmol) was dissolved in 0.5 mL of DMSO. Finally, these two reactions mixtures were mixed and stirred for 24 hours at room temperature. After 24 hours, the reaction mixture was dialyzed against water for 6-8 hours using dialysis membrane with MWCO 3.5KDa. This dialyzed reaction mixture was then frozen and lyophilized to obtain a solid powder. This powder was washed several times with methanol acetone solvent mixture and the obtained solid product was separated by centrifugation. Thereafter, this solid product was dissolved in water, frozen and lyophilized to obtain a white powder as a pure product (**Figure 2**, **Figure S2e**). Yield = 0.55 mg (78 %). $^1$H-NMR (500 MHz, $d_6$-DMSO) δ ppm: : 7.1-7.5 (4H, AZD5069), 6.1(1H, AZD 5069), 5.5-3.0 (broad peaks, dextran hydroxyl, glucosidic and PEG protons ) 4.63 ppm (s, dextran anomeric proton), 2.6 ppm (m,4H) methylene protons of SA, 1.48 ppm, 1.25-0.85 ppm (aliphatic protons for stearic acid and AZD5069).

***1.6. Biophysical characterization of CXCR2-homing nanoparticle.***
For the 1H-NMR studies, the samples with appropriate solvents were loaded onto NMR tubes and the proton NMR spectra were recorded using 500MHz Varian Spectrophotometer. The Dynamic Light Scattering measurements and Zeta potential were captured using Malvern Zetasizer Nano ZS instrument. The conjugated nanoparticles were resuspended in strerile PBS and analyzed for size and surface potential at different timepoints. High Performance Liquid Chromatography (HPLC) analysis were performed on Agilent 1200 series equipped with UV-visible and florescent detectors and automated injectors. TEM images were acquired using Philips/FEI Tecnai 20 microscope.

***1.7. Single-cell RNA sequencing (scRNAseq) analysis.*** ScRNAseq analysis from *Ptf1a*$^{cre/+}$;*LSL-Kras*$^{G12D/+}$;*Tgfbr2*$^{flox/flox}$ (PKT) tumors was performed as described previously[1]. The 10x Genomics Chromium Single Cell 3' Reagent v3.1(Cat # PN-1000268) was used with standard conditions and volumes to process cell suspensions for 3' transcriptional profiling. Single cell suspensions from PKT mice were extracted, and live cells were sorted using flow cytometry, and volumes were calculated for a target cell recovery of 100,000 cells and loaded on the Chromium Controller as per manufacturer guidelines. The resultant purified cDNA was quantified and qualitative assessed on the Agilent Bioanalyzer using the High Sensitivity DNA Kit (Cat #5067-4626). The final single-cell 3' libraries were quantified using the Qubit dsDNA High Sensitivity (Cat #Q33231) and qualitatively evaluated on the Agilent Bioanalyzer using the High Sensitivity DNA Kit. For sequencing, libraries were loaded at optimized concentrations onto an Illumina NovaSeq and paired end sequenced under recommended settings (R1: 28 cycles; i7 index: 10 cycles; i5 index: 10 cycles; R2: 90 cycles). The libraries were diluted to varying nM concentrations in Illumina Resuspension Buffer (PN-15026770), denatured according to Illumina standard guidelines, and loaded on the Illumina NovaSeq at 1.2 nanomolar. The resulting intensity files were demultiplexed as FASTQ files using Illumina BaseSpace software and then aligned to the transcriptome using the 10x Genomics CellRanger (ver4.0.0) software package. The 16 human PDAC scRNAseq dataset by Steele et al.[2] were retrieved from GEO with the access code GSE155698 and processes accordingly to their code: https://github.com/PascaDiMagliano-Lab/MultimodalMappingPDA-scRNASeq.

***1.8. Cluster identification and gene annotation in scRNAseq dataset*.** Principal component analysis was performed on the scaled data to reduce the dimensions, with number of components chosen based on a cumulative proportion (accumulated amount of explained variance) of 90%. Cells with >5% mitochondrial counts, less than 200 or more than 2500 unique feature counts were filtered out. Clusters of cells were identified by shared nearest neighbor algorithm. Cell type annotations were assigned to clusters based on the expression of canonical features in a minimum percentage of cells. Next, the function “FindMarkers” from Seurat v4.0 R package was utilized to identify differentially expressed genes for identity classes. Genes were considered differentially expressed if detected in at least 25% of clusters, with default log(fold change) of 0.25.

***1.9. Cell Lines.*** J774 cells have been shown to phenocopy PMN-MDSCs *in vitro*[3]. Culture conditions for J774M cells were as follows: 37°C, 5% CO2 in RPMI 1640 supplemented with 10% FBS, 1.5% HEPES buffer, 1% L-glutamine, 1% MEM non-essential amino acids, 1% penicillin-streptomycin, 1% sodium pyruvate, 0.0004% beta-mercaptoethanol (Sigma Aldrich). All cell lines were stored in liquid nitrogen and frozen between 5-10 passages from the original cell line. No cell lines were used beyond 15 passages. Cell lines were regularly tested for Mycoplasma every three months in accordance with laboratory policy.

***1.10. Arginase Activity Luminescence Assay*.** Assay was performed according to manufacturer protocol. In particular, cells were lysed with Arginase Assay lysis buffer at $1x10^6$ cells/100µL and plated at $5x10^5$ cells/well in two flat-bottom wells of a low-retention plate per replicate, carefully avoiding bubble formation, and topped with 40µL of Assay Buffer. Target samples were incubated for 20 minutes at 37°C with 10µL of included $H_2O_2$ substrate solution, while background wells were incubated with an additional 10µL of buffer. Standards were prepared and plated in duplicates per kit instructions, followed by the enzymatic reaction mixture which was added to all wells. Raw absorbance values were immediately obtained every 2 minutes over a 30-minute period using a plate reader (Molecular Devices SpectraMax M3) at OD=570 nm at 37°C. Arginase Activity Units were then calculated from raw absorbance values. ΔOD (ΔOD= (OD2-ODbg2)-(OD1-ODbg1)) was used to obtain the nmol of $H_2O_2$ generated by arginase, collected from a standard curve of known $H_2O_2$ concentrations. Arginase activity was calculated as $(B/\Delta T*V)*D$ in units/mL, where B is amount of $H_2O_2$ from standard curve (nmol), V is the sample volume added into reaction well (mL), and D is sample dilution factor. One unit of Arginase activity refers to the amount of arginase that will generate 1.0 nmol of $H_2O_2$ per minute at pH 8 at 37°C.

***1.11. Enzyme-linked immunosorbent assay (ELISA).*** Condition media was quantified, and equal amount of protein loaded per well of a standard 96 well plate, including three technical replicates per biological replicate. ELISA kits specific for IFN-γ (R&D Systems) were used according to the manufacturer's protocol.

***1.12. Western Blotting*.** Single cell suspensions were homogenized (Omni™ Tissue Homogenizer) in 0.5 ml of RIPA lysis buffer (20-188, Millipore®) and sonicated for 2 minutes on ice. Tissue/cell lysates were spined for 10’ at 15000 rpm at 4°C. Supernatants were then collected, quantified with BCA assay (23227; ThermoFisher Scientific). Lysates containing 30-40 µg of equal protein were separated using 4-20 % SDS PAGE Mini-PROTEAN TGX Stain-Free Gel (Bio-Rad) and transferred on Trans-Blot® Turbo™ Midi PVDF Transfer Packs (Bio-Rad) using Trans-Blot Turbo Transfer System (Bio-Rad). For immunodetection, membranes were incubated with primary antibodies listed in **Table S1** at 4°C overnight. Membranes were then washed and incubated with corresponding specific secondary antibodies conjugated with horseradish peroxidase (Jackson Immuno Research Laboratory). Immunoreactive bands were developed using Pierce ECL Western Blotting Substrate (Thermo Scientific) or SuperSignal™ West Pico PLUS Chemiluminescent Substrate (Thermo Scientific). Densitometric analysis for quantification of protein expression was performed using ImageJ software.

***1.13. Quantitative Polymerase Chain Reaction (qPCR).*** Purified RNA was obtained from in vitro cells or tumor samples using RNeasy Kit (Qiagen) according to manufacturer's protocol. RNA concentration and quality were verified using NanoDrop spectrophotometer (ThermoFisher). cDNA was then generated by reverse transcription of RNA product using High-Capacity cDNA Reverse Transcription Kit with RNase Inhibitor (Applied Biosystems). qPCR was performed using incubation with gene specific predesigned arginase-1 (RT2 qPCR Primer Assay, Qiagen) to gene targets and iQTM SYBR® Green Supermix (BioRad). Gene expression was normalized to the housekeeping gene GAPDH 0r 18S using ΔΔCT method and reported as fold change relative to control.

***1.14. Orthotopic and subcutaneous mouse models.*** *Orthotopic model.* These were performed as described previously (1,6,7) .C57BL/6 mice were anesthetized with ketamine/xylazine (10:1) in sterile saline and, under sterile conditions, KPC6694c2 tumor cell suspensions (ranging from $25x10^3$-$1x10^5$ cells/injection) in Matrigel® (Corning, #354230) were injected directly into the pancreata of mice with subsequent closure of the peritoneum with Vicryl 5-0® sutures and skin with skin staples. Tumors were allowed to grow for 1-2 weeks prior to treatment initiation.

*Subcutaneous model.* The flank injection model was generated by subcutaneously injecting $5x10^5$ KPC tumor cells resuspended in Matrigel® (Corning, #354230) 50% in saline in the flanks of C57BL/6 mice. The animals were monitored and sacrificed prior to tumors reaching a total volume>$10^3$ $mm^3$.

***1.15. Tissue Processing*.** Whole pancreata harvested from KPC tumor-bearing mice were enzymatically digested utilizing a solution comprising 0.6 mg/ml collagenase P (Sigma Aldrich, #11213857001), 0.8 mg/ml Collagenase V (Sigma Aldrich, #9001-12-1), 0.6 mg/ml soybean trypsin inhibitor (Sigma Aldrich, # 9035-81-8), and 1800 U/ml DNase I (Thermo Fisher Scientific, #18047019) in RPMI medium for 20-30 minutes at 37°C. Samples were then washed and underwent single-cell dissociation using 40 μm smash strainers. and single-cell suspensions were processed using RBC lysis buffer. Samples were then kept frozen at -80°C.

***1.16. Flow Cytometry***. Single cell suspensions from mouse tumors or patient derived peripheral blood cells were thawed, washed with autoMACS rinsing buffer + BSA (Miltenyi Biotec, #130-091-222, #130-091-376), and incubated with FcR-blocking reagent (Miltenyi Biotec, #130-092-575), and subsequently stained with fluorescently conjugated antibodies (**Table S2**). Ghost Red Dye 780 (TONBO biosciences, #13-0865) or LIVE/DEAD™ Fixable Blue (Invitrogen, #L23105) live/dead discrimination was performed as per manufacturer's protocol and cells were fixed with 1% formaldehyde solution (Thermo Fisher Scientific, #047392-9M). Flow cytometry data acquisition was performed on Cytek Aurora and analyzed using FlowJo v.10 software. For viSNE plot generation, flow cytometry data were analyzed using FlowJo v.10 with default parameters (iterations = 1000, perplexity = 30, θ = 0.5). All samples were derived from the same viSNE run by combining all individual flow cytometry standard files into a single flow cytometry standard file using the concatenation tool.

***1.17. IVIS imaging experiments.*** Tumor or non-tumor bearing (flank/subcutaneous) C57BL/6 mice were injected using either intraperitoneal or intravenous routes with (1) PBS; (2) non dye-containing CXCR2-NP; (3) non-AZD5069 decorated NP encapsulating Cy5.5 dye; (4) CXCR2-$NP^{Cy5.5}$ in non tumor-bearing mouse; and (5) CXCR2-$NP^{Cy5.5}$. 24 hours following injection, mice were temporarily anesthetized using isoflurane and imaged using IVIS Spectrum optical imaging system (PerkinElmer).

***1.18. T-cell suppression assays*.** J774 cells pre-conditioned with either CXCR2-$NP^{Ruxo}$ or free Ruxolitinib drug were co-cultured with CD3/28-stimulated $CD3^{+}$ T-cells and T-cell activity was measured via IFN-γ secretion. T-cells were isolated from spleens of tumor-naïve C57BL/6 mice using magnetic column PanT cell isolation kit as per manufacturer's recommendation (Miltenyi Biotec, #130-095-130). T-cells were co-cultured with J774 cells at 1:2 T-cell:J774) and anti-CD3/CD28 beads (Thermo Fisher Scientific) at a bead-to-T-cell ratio of 1:1. After 48h and 72h, condition media was collected for IFN-γ measurement by ELISA.

### 2. SUPPLEMENTARY TABLE

#### 2.1. Table S1. Western Blotting Antibodies

| Marker | Species | Company | Catalog # |
|---|---|---|---|
| Total STAT3 | Rabbit | Cell Signaling | 12640S |
| p-$STAT3^{Y705}$ | Rabbit | Cell Signaling | 9145S |
| Vinculin | Rabbit | Cell Signaling | 13991S |
| β-Actin | Mouse | Cell Signaling | 3700S |

#### 2.2. Table S2. Flow Cytometry Antibodies

| **Myeloid Markers** | | | |
|---|---|---|---|
| **Target** | **Fluorophore** | **Catalog #** | **Company** |
| CD45 | FITC | 157608 | Biolegend |
| CD11b | AF700 | 101222 | Biolegend |
| F4/80 | BV785 | 123141 | Biolegend |
| CD86 | BV421 | 105123 | Biolegend |
| MHC II | PEDazzle | 107648 | Biolegend |
| CD206 | PE-Cy7 | 141720 | Biolegend |
| LY6G | BV650 | 108441 | Biolegend |
| LY6C | PerCP-Cy™5.5 | 128012 | Biolegend |

| | | | |
|---|---|---|---|
| CXCR2 | APC | 149312 | Biolegend |
| **Lymphocyte Markers** | | | |
| **Target** | **Fluorophore** | **Catalog #** | **Company** |
| TCRb | BUV496 | 749915 | BD |
| CD62L | BUV563 | 741230 | BD |
| CD44 | BUV737 | 612799 | BD |
| CD45 | BUV805 | 741957 | BD |
| CD8a | BV650 | 301042 | Biolegend |
| IFN-γ | PE | 505807 | Biolegend |
| CD4 | BV785 | 100552 | Biolegend |
| PD-1 | PE-Dazzle 594 | 121624 | Biolegend |

### 3. SUPPLEMENTARY FIGURES

**Figure S1:** Gene expression of *Arg1* from scRNAseq analysis of cells dissociated from tumors in PKT genetically engineered mice. UMAP projection separates various intratumoral cell types as described previously[4].

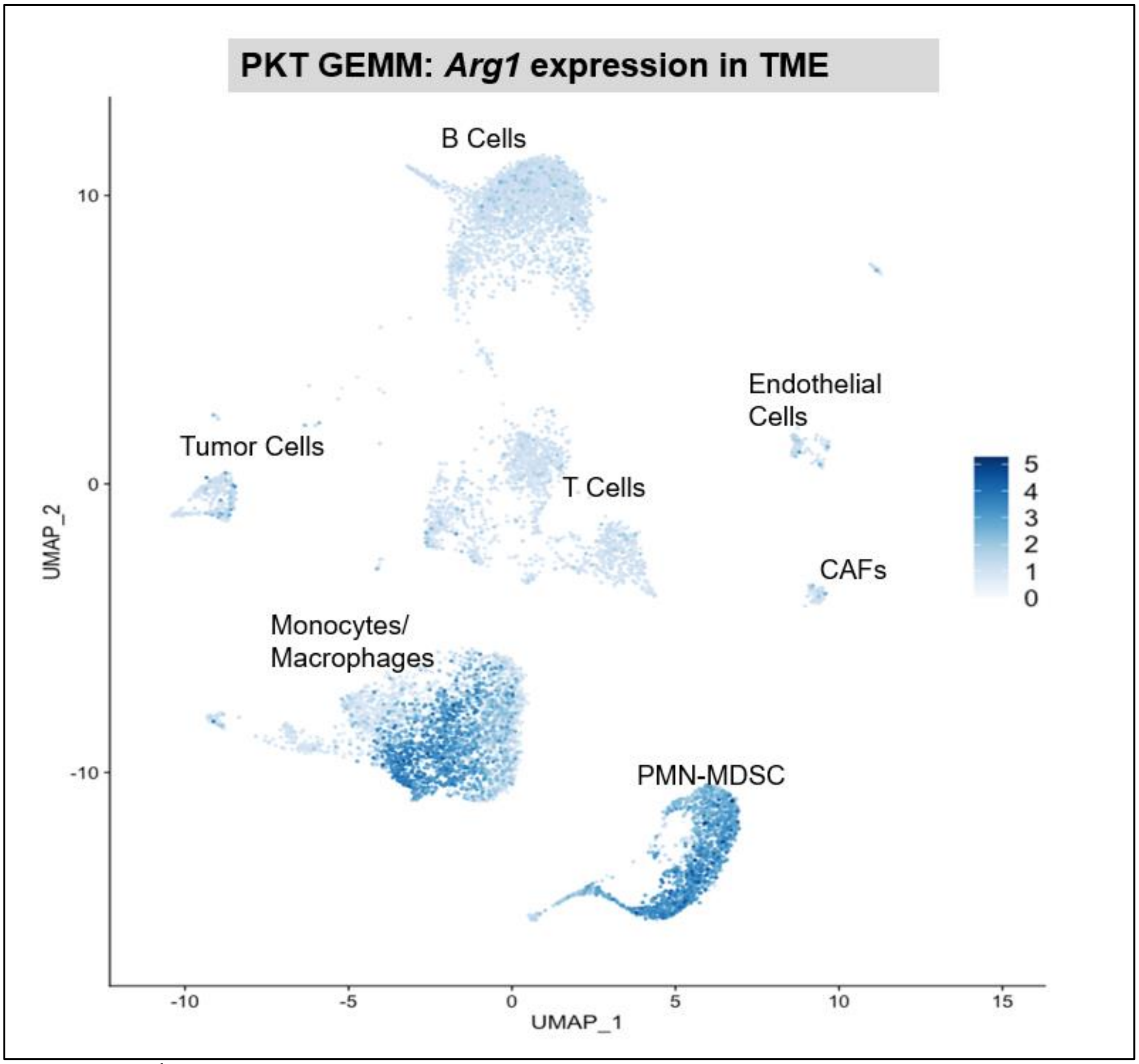


**Figure S2:** [1]H-NMR spectra validating intermediate products of nanoparticle synthesis, recorded on a 400 and 500 MHz Bruker NMR instrument. **a,** Dextran-stearic acid-succinic mono tert. butyl ester derivative [1]H-NMR (400 MHz, $d_6$-DMSO) δ ppm: 4.47, 4.82, 4.88 ppm (s, hydroxyl of dextran) 4.63 ppm (s, dextran anomeric proton), 3.14-3.69 ppm (dextran glucosidic protons), 2.6 ppm (m,4H) methylene protons of SA, 1.48 ppm,1.39 ppm (s, 9H) tert. butyl, 1.25-0.85 ppm (aliphatic protons). **b,** Dextran-stearic acid-succinic acid derivative [1]H-NMR (500 MHz, $d_6$-DMSO) δ ppm: 4.47, 4.82, 4.88 ppm (s, hydroxyl of dextran) 4.63 ppm (s, dextran anomeric proton), 3.14-3.69 ppm (dextran glucosidic protons), 2.6 ppm (m,4H) methylene protons of SA, 1.48 ppm, 1.25-0.85 ppm (aliphatic protons). **c**, Boc-NH-PEG-AZD5069. **d,** $NH_2$-$(PEG)_{36}$-CO-AZD5069 Conjugate [1]H-NMR (500 MHz, $d_6$-DMSO) δ ppm: 7.1-7.5 (4H, AZD5069), 6.1(1H, AZD 5069), 4.44 (2H, PEG), 3.3-3.8 (165H, PEG and AZD5069), 3.0-3.1 (8H, PEG + AZD 5069), 1.18-1.22 (5H, from AZD5069). **e**, Dex-Stearic-PEG-AZD5069.

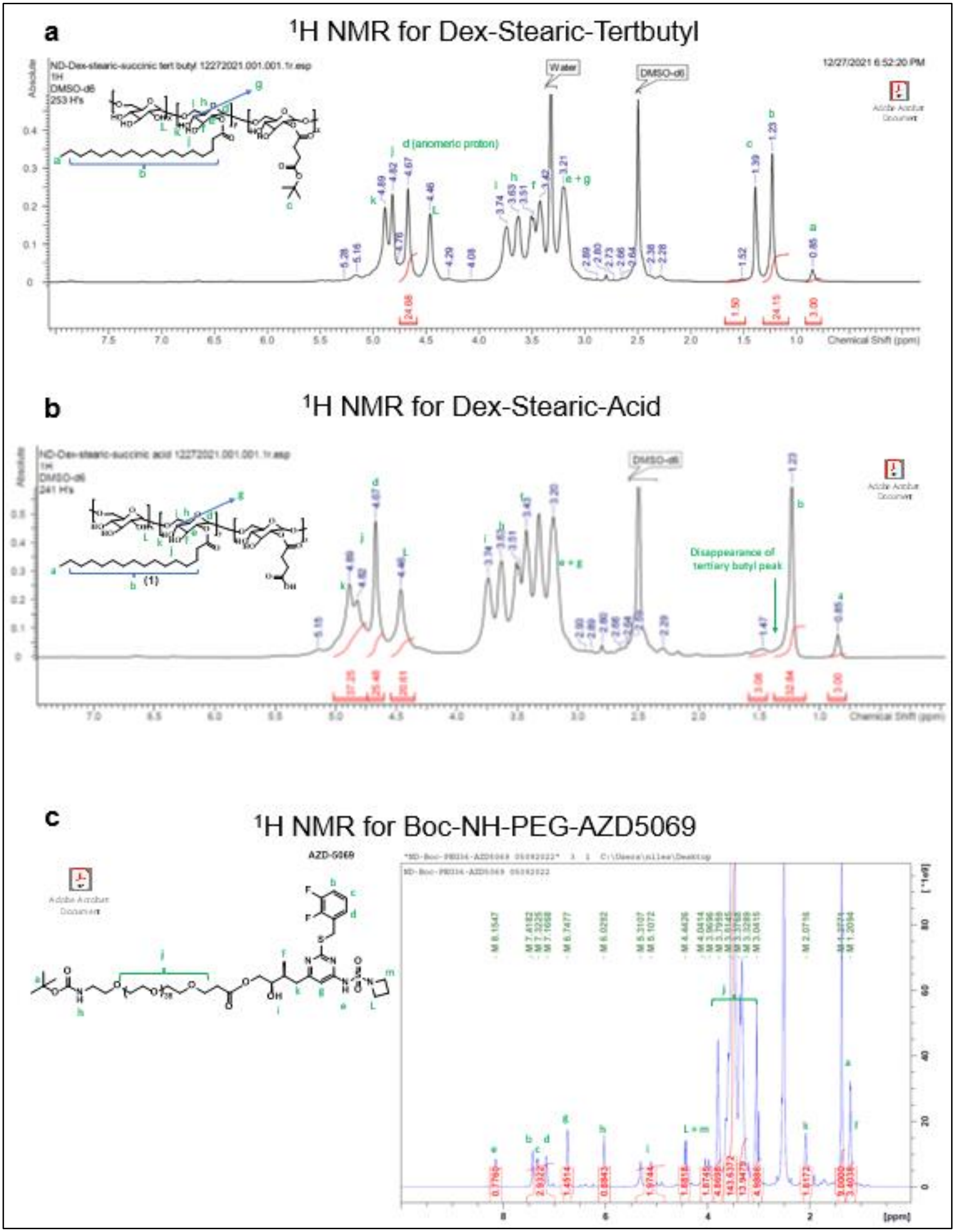
a
1H NMR for Dex-Stearic-Tertbutyl
ND-Dex-stearic-succinic tert butyl 12272021.001.001.1r.esp
1H
DMSO-d6
253 H's
12/27/2021 6:52:20 PM
Water
DMSO-d6
d (anomeric proton)
e + g
Chemical Shift (ppm)
b
1H NMR for Dex-Stearic-Acid
DMSO-d6
241 H's
Disappearance of tertiary butyl peak
Chemical Shift (ppm)
c
1H NMR for Boc-NH-PEG-AZD5069
AZD-5069
L + m
[ppm]

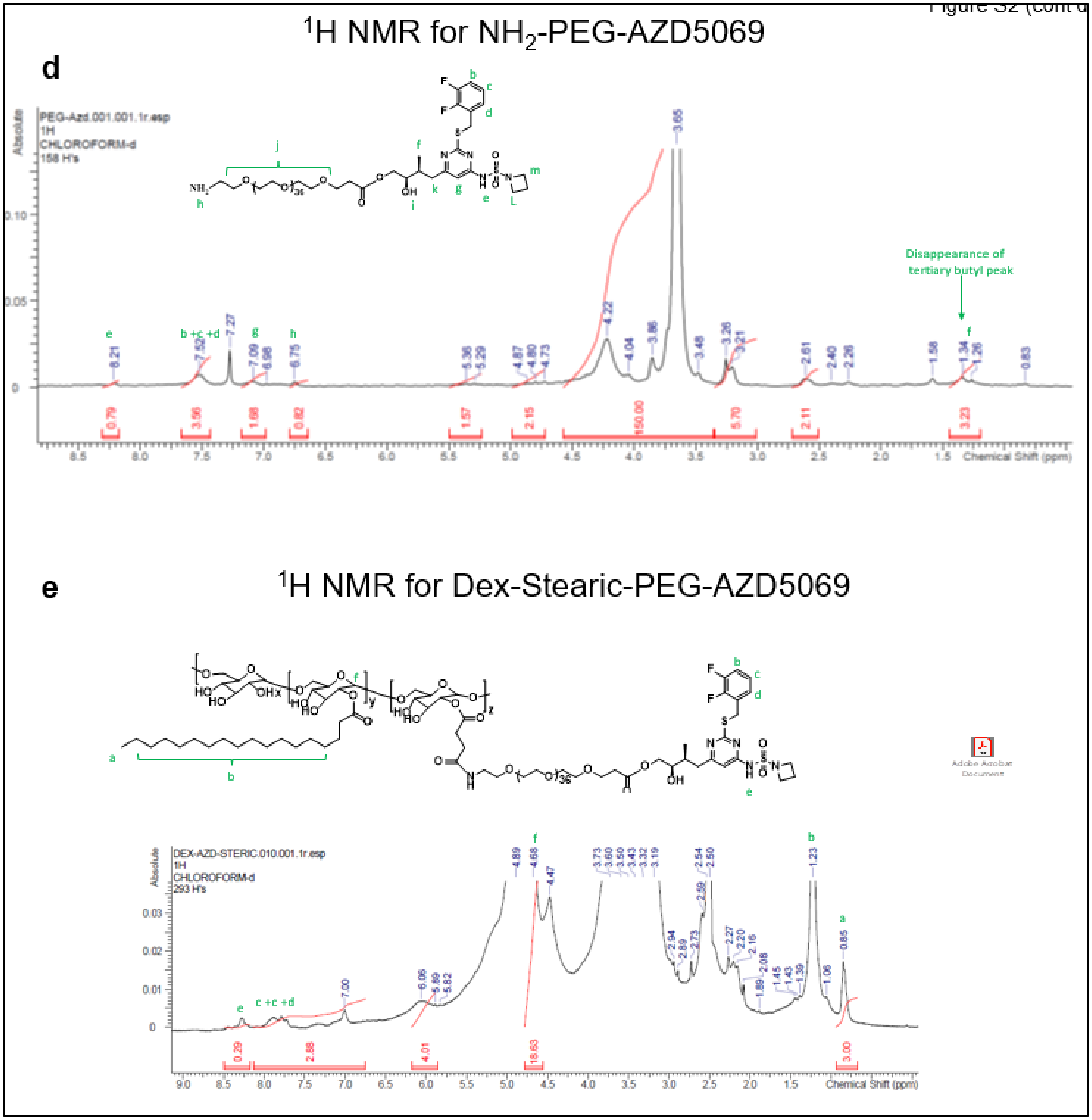
[illegible]
d
$^1$H NMR for NH$_2$-PEG-AZD5069
PEG-Azd.001.001.1r.esp
1H
CHLOROFORM-d
158 H's
Disappearance of tertiary butyl peak
Chemical Shift (ppm)
Absolute
e
$^1$H NMR for Dex-Stearic-PEG-AZD5069
DEX-AZD-STERIC.010.001.1r.esp
1H
CHLOROFORM-d
293 H's
Adobe Acrobat Document
Chemical Shift (ppm)
Absolute

**Figure S3: a**, Flow cytometry gating strategy showing how J774 cells phenocopy CXCR2⁺ PMN-MDSCs *in vitro*. **b,** CCK-8 assay assessing cell viability of J774 cells with serial concentrations of non-decorated nanoparticles and CXCR2-NP, both encapsulating Cy5.5 dye, at 72 hours. 10μL of CCK-8 reagent was added to each well and absorbance at 450 nm was measured using a microplate reader after 4 hours of incubation. The cell viability was calculated by comparing the absorbance of each well to that of a control well and plotted as percentage (n=3 per concentration).

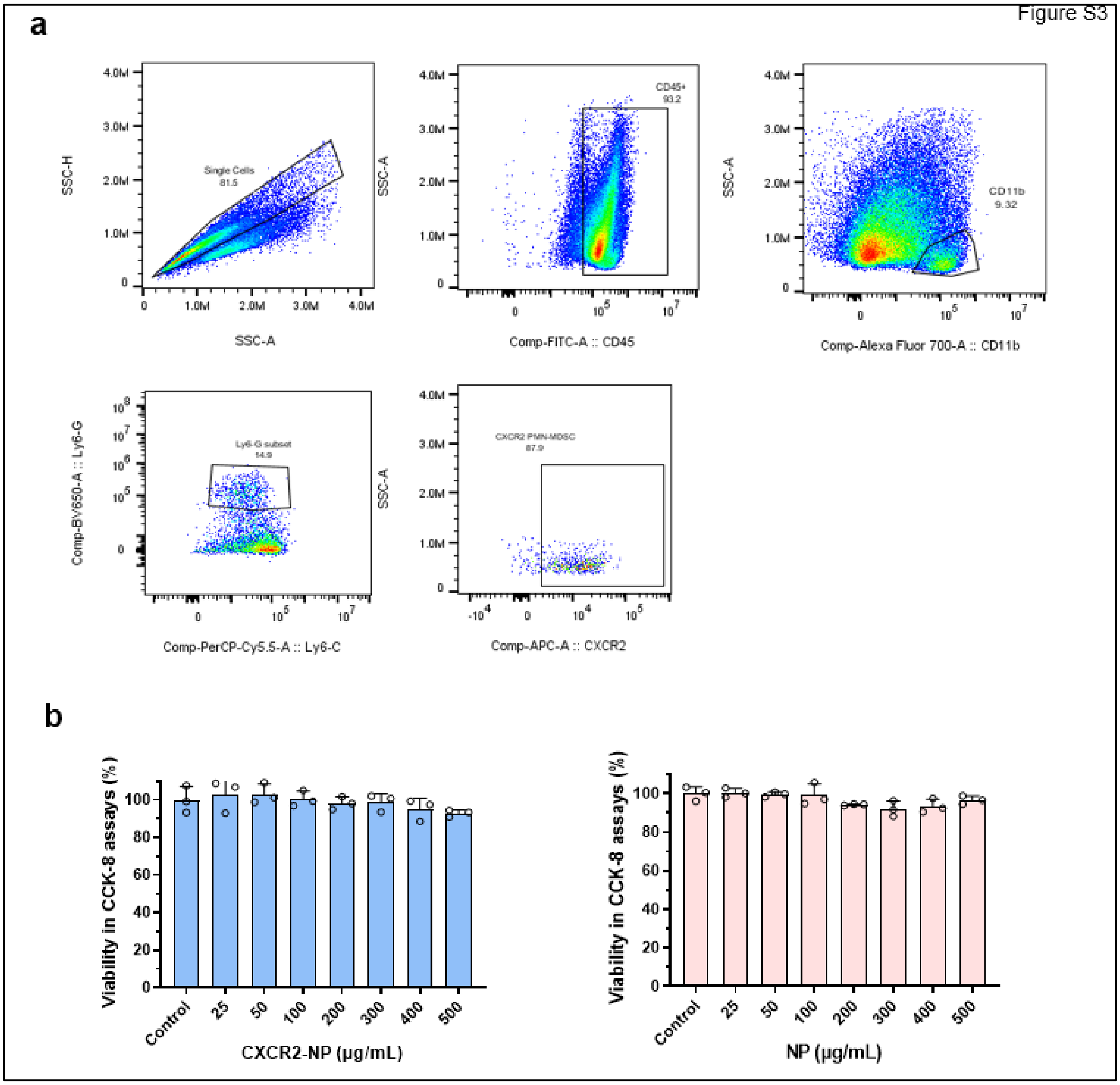

**Figure S4: a,** Quantification of cells staining positive for Cy5.5 dye fluorescence were measured utilizing Image J software (n=3 per group). **b,** Receptor mediated uptake of CXCR2-NP$^{Cy5.5}$ nanoparticles in different cell types within tumor microenvironment at 24 hours, visualized and quantified by confocal microscopy. Red fluorescence from Cy5.5 was monitored through the red channel (λ = 594 nm), green fluorescence from phalloidin was monitored through a green channel (λ = 488 nm) and the blue fluorescence produced by cell nuclei hoechst staining was monitored through a blue channel (λ = 405). All the confocal images were recorded keeping all the parameters constant for every measurement.

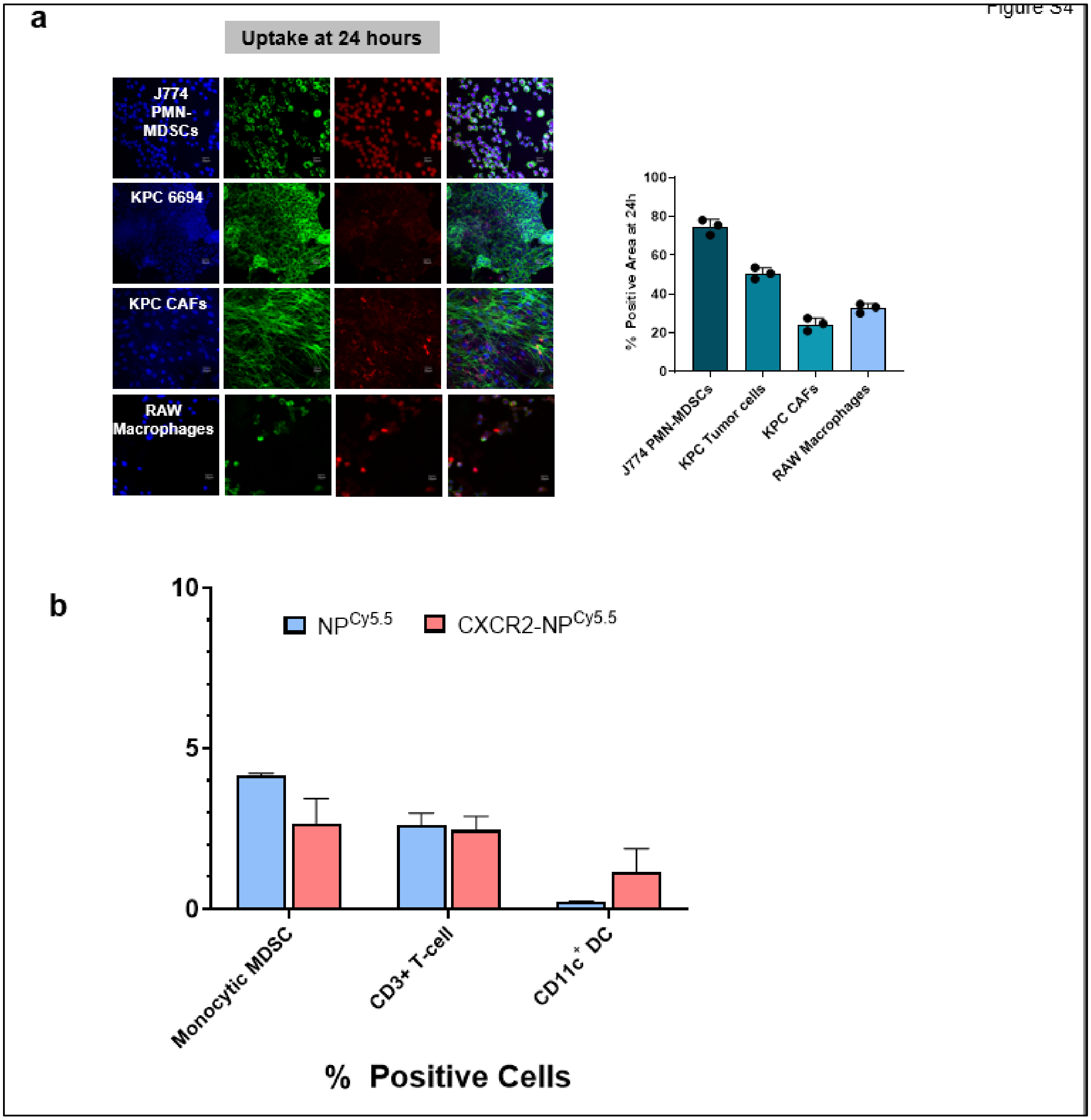

**Figure S5: a,** Viability of neutrophils in spleens of mice in respective treatment cohorts at endpoint analysis (day 30) was quantified via flow cytometry and plotted as a frequency of F4/80$^-$CD11b$^+$ cells (n=3 mice/group). **b**, Absolute numbers of intratumoral CD4$^+$ T-cells gated on CD3$^+$ cells from mice in indicated treatment groups, and compared using one-way ANOVA.

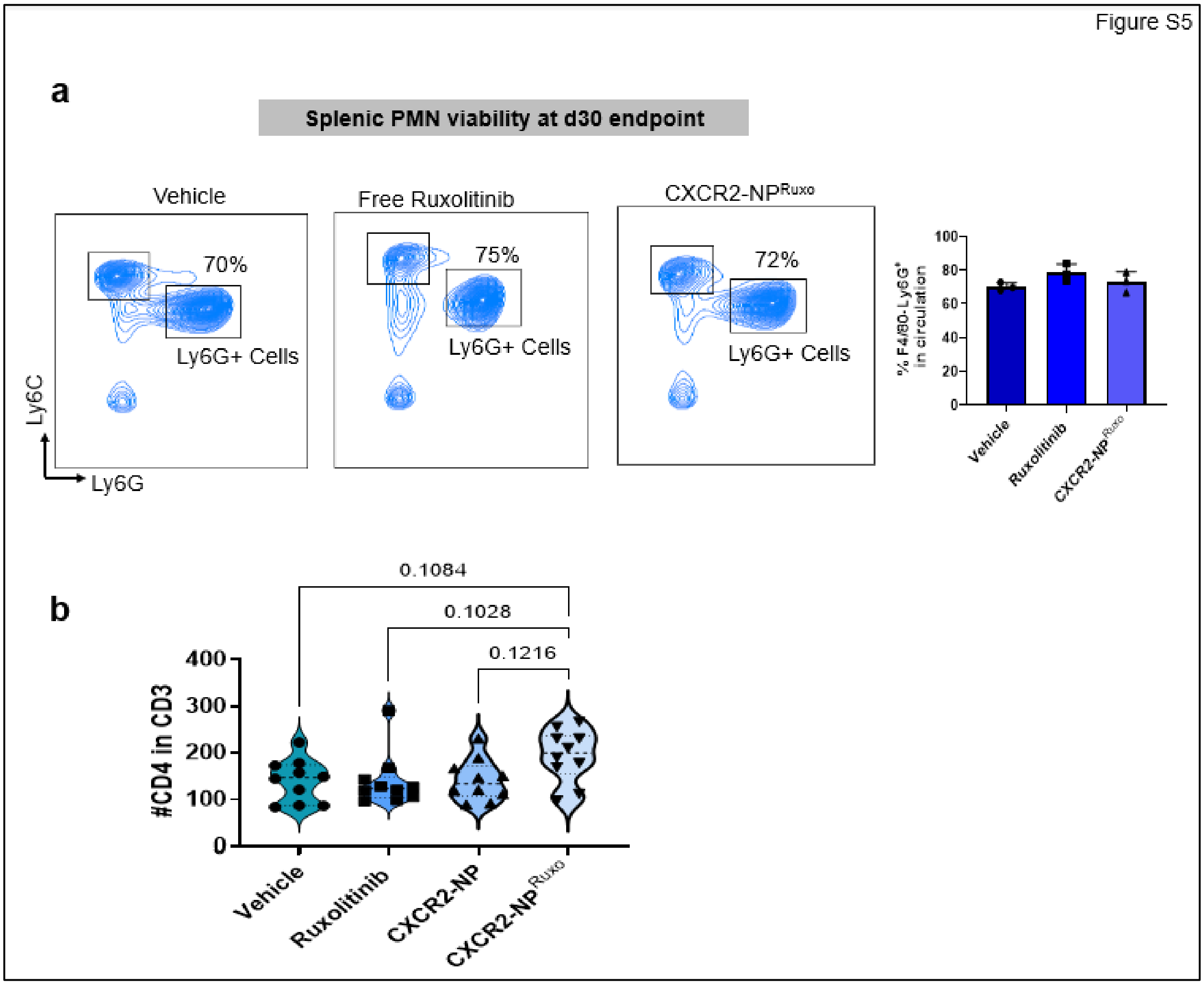
